# Social belonging: Brain structure and function is linked to membership in sports teams, religious groups and social clubs

**DOI:** 10.1101/2021.09.06.459167

**Authors:** Carolin Kieckhaefer, Leonhard Schilbach, Danilo Bzdok

## Abstract

Human behaviour across the life span is driven by the psychological need to belong, from kindergarten to bingo nights. Being part of social groups constitutes a backbone for communal life, and confers many benefits for physical and mental health. Capitalizing on neuroimaging and behavioural data from ~40.000 participants from the UK Biobank population cohort, we used structural and functional analyses to explore how social participation is reflected in the human brain. Across three different types of social groups, structural analyses point towards variance in ventromedial prefrontal cortex, fusiform gyrus and anterior cingulate cortex as structural substrates tightly linked to social participation. Functional connectivity analyses emphasized the importance of default mode and limbic network, but also showed differences for sports teams and religious groups as compared to social clubs.

Taken together, our findings establish the structural and functional integrity of the default mode network as a neural signature of social belonging.

## Introduction

Belonging to social groups is an indispensable ingredient of everyday life, providing well-documented benefits for mental health and well-being. Regular engagement in social groups of varying complexity has been structuring societies for thousands of years. Experiencing the team spirit in a vibrant soccer match, enjoying a beer in a bar with like-minded persons or resting in collective silent prayer - these are formative experiences that scaffold human social life. These group events are some of many examples of social participation.

The definition of social participation has taken many forms in previous work. Yet, a content analysis revealed the “involvement in activities that provide interactions with others” as a key recurring theme (Levasseur, Richard, et al., 2010). In contrast, the type of activity or level of involvement with the group are more variable. Social participation is associated with enhanced subjective and objective health outcomes (Leone & Hessel, 2016; Sirven & Debrand, 2008), reduction of depressive symptoms (Chiao et al., 2011) and improved overall quality of life (Lestari et al., 2021; Levasseur, Desrosiers, et al., 2010). A representative population-based study in the US showed lower levels of self-rated physical health for people who experience social disconnectedness and perceived isolation (Cornwell & Waite, 2009). The central relevance of social participation was also underscored by the WHO in the context of aging and rehabilitation (WHO, 2002), finding its way into political programs and recommendations aiming to improve global health in disadvantaged populations (Francés et al., 2016). The recent European health policy framework *Health 2020* emphasized participation as a core principle for health equity and well-being (Boyce & Brown, 2017).

The close links between social participation and well-being holds across the life span and is not limited to aging. A transnational study investigated 1047 participants with an average age of 35.5 years from Europe, North-Africa, Western Asia and the Americas in terms of mental health in a period of pandemic-related restrictions on social relations (Ammar et al., 2020). A decrease of social participation due to these restrictions took a negative toll on life satisfaction. Comparing Italian, American and Iranian students, a different study showed positive associations between social participation and well-being, largely mediated by the sense of community (Cicognani et al., 2008). The concept of social participation is closely related to sense of community, sharing several positive effects on individual and social life (Talò et al., 2014).

The sense of community emphasizes the aspect of shared emotional connection of a membership in a community or group. This aspect might be an important explanatory factor of the high social gain in relationships within groups in the periphery of one’s social network. Resulting feelings of connectedness with others in conjunction with social participation was recently discussed as a buffer for less decline in well-being in the sense of a resource (Sharifian & Grühn, 2019). Belonging to a group helps to increase self-acceptance, thus being an important factor in coping processes, especially in challenging situations (Thoits, 2011). Furthermore, the sense of belonging directly and persistently contributes to people’s resilience (Scarf et al., 2016).

A further concept that is closely linked to resilience is the sense of meaning and purpose in life. It denotes the skills and readiness for mastering difficult circumstances (Feldman, 2020). A recent study links the association between social interaction and meaning in life to the brain (Mwilambwe-Tshilobo et al., 2019). These authors reported an increased functional connectivity between the default and limbic networks to be associated with a greater sense of life meaning. The increase of perceived meaning in life was repeatedly associated with social participation and belonging to others (Adams et al., 2011; Sharifian & Grühn, 2019). Hence, current limitations of social contacts are a challenge for purposeful living.

In the context of the COVID-19 pandemic with widespread loneliness and reduced possibilities of participation due external circumstances, individual resilience gains relevance from a public health standpoint. Being able to cope with and quickly recover from a disaster is critically dependent on active belonging to a group within the resilience process (Quinn et al., 2020). A closer look at the biological manifestations of social belonging and participation at the population-level is imperative, especially as COVID-19 challenged not only regions or communities but affected entire populations. Such confronted life events and overcome obstacles are important to be investigated for the angle of psychopathology and clinical symptoms.

In the present population-based approach, we shift the focus towards factors that are known to be key for successful resilience and mental well-being. Regarding how a particular individual handles stressors, previous studies underlined the association between the experience of control as well as resilient coping and the activation of the medial prefrontal cortex (Maier & Watkins, 2010; Sinha et al., 2016). Neuroimaging data from real-life contexts offers important insight into social belonging and its many wide-ranging consequences. One way to understand the broader link between social interactions and its influence on brain architecture has been proposed by the social brain hypothesis. The intensified need and intricacy of social relationships in humans may have spurred refinement towards more complex representation of social bonds in the brain (Dunbar, 2009). Brain systems that have long been described to be closely implicated in social cognition processes involve the default mode network (DMN; Mars et al., 2012). However, based on previous works, the neural basis of social participation and belonging remain obscure. Taken together no human brain-imaging assessments of participation in different group contexts exist, in part because such characteristics of people’s everyday social life have seldom been systematically acquired before the emergence of large population datasets such as the UK Biobank cohort. Combining structural and functional neuroimaging as well as demographic profiling with a population-based approach, we investigated three particular forms of social participation: sports teams, religious groups and social clubs. Using a population cohort to examine and understand systematic variations of social belonging and resilience help inform public-health decision making, which ultimately can foster implementation and even interventions in practice.

## Materials and Methods

### Data resources

The UK Biobank is a prospective epidemiology resource that offers extensive behavioural and demographic assessments, medical and cognitive measures, as well as biological samples in a cohort of ~500,000 participants recruited from across Great Britain (https://www.ukbiobank.ac.uk/). This openly accessible population dataset aims to provide multimodal brain-imaging for ~100,000 individuals, planned for completion in 2022. The present study was based on the recent data release from February/March 2020 that augmented brain scanning information to ~40,000 participants.

In an attempt to improve comparability and reproducibility, our study built on the uniform data preprocessing pipelines designed and carried out by FMRIB, Oxford University, UK (Alfaro-Almagro et al., 2018). We involved data from the ~40,000 participant release with brain-imaging measures of grey matter morphology (T1-weighted MRI [sMRI]) and neural activity fluctuations (resting-state functional MRI [fMRI]) from 48% men and 52% women, aged 40-69 years when recruited (mean age 54.9, standard deviation [SD] 7.5 years). Our study focused on regular social engagement as captured by membership in social group (Bzdok & Dunbar, 2020; Hawkley et al., 2003; Luhmann & Hawkley, 2016). This self-reported item was based on the following question: “Which of the following do you attend once a week or more often?” (data field 6160). Our study focussed on three target groups: people reporting engagement in sports teams, religious groups and social clubs.

Similar measures are found in widely used assessments of social embeddedness (Cohen & Hoberman, 1983; Cyranowski et al., 2013; Hawkley et al., 2005). Conceptually similar (Cyranowski et al. 2013) are also contained in other standard measurement-tools of social embeddedness, such as the Revised UCLA Loneliness Scale (Hawkley et al. 2005) and the Interpersonal Support Evaluation List (Cohen and Hoberman 1983). A variety of studies showed single-item measures of social traits to be reliable and valid (Atroszko et al., 2015; Dollinger & Malmquist, 2009; Mashek et al., 2007). Previous research has used such individual items for successfully measuring social support (Atroszko et al. 2015), community connectedness (Mashek et al. 2007), and perceived social isolation (Ong et al., 2016). For example, the separate item “There are people I can talk to” correlates very highly (r = .88) with the specific dimension of the R-UCLA Loneliness Scale that corresponds to the objective frequency of social interaction (Hawkley et al. 2005).

The present analyses were conducted under UK Biobank application number 25163. All participants provided informed consent. Further information on the consent procedure can be found elsewhere (http://biobank.ctsu.ox.ac.uk/crystal/field.cgi?id=200).

### Multimodal brain-imaging and preprocessing procedures

Magnetic resonance imaging scanners (3T MRI Siemens Skyra) were matched at several dedicated imaging sites with the same acquisition protocols and standard Siemens 32-channel radiofrequency receiver head coils. To protect the anonymity of the study participants, brain-imaging data were defaced and any sensitive meta-information was removed. Automated processing and quality control pipelines were deployed (Alfaro-Almagro et al., 2018; Miller et al., 2016). To improve homogeneity of the imaging data, noise was removed by means of 190 sensitivity features. This approach allowed for the reliable identification and exclusion of problematic brain scans, such as due to excessive head motion.

#### Structural MRI

The sMRI data were acquired as high-resolution T1-weighted images of brain anatomy using a 3D MPRAGE sequence at 1 mm isotropic resolution. Preprocessing included gradient distortion correction (GDC), field of view reduction using the Brain Extraction Tool (Smith, 2002) and FLIRT (Jenkinson et al., 2002; Jenkinson & Smith, 2001), as well as non-linear registration to MNI152 standard space at 1 mm resolution using FNIRT (Andersson et al., 2007). To avoid unnecessary interpolation, all image transformations were estimated, combined and applied by a single interpolation step. Tissue-type segmentation into cerebrospinal fluid (CSF), grey matter (GM) and white matter (WM) was applied using FAST (FMRIB’s Automated Segmentation Tool, (Zhang et al., 2001) to generate full bias-field-corrected images. SIENAX (Smith et al., 2002), in turn, was used to derive volumetric measures normalized for head sizes.

#### Functional MRI

The fMRI data of intrinsic neural activity were acquired without engagement in a predefined experimental task context at 2.4 mm spatial resolution, time to repeat=0.735s, and with multiband acceleration of 8. A single-band reference image with higher between-tissue contrast and without T1-saturation effects was acquired within the same geometry as the time series of neural activity maps. The reference scan was used for the alignment to other brain-imaging modalities and correction for head motion. Preprocessing was performed using MELODIC (Beckmann & Smith, 2004), including EPI and GDC unwarping, motion correction, grand-mean intensity normalization, and high-pass temporal filtering (Gaussian-weighted least-squares straight line fitting, sigma=50s). The ensuing images were submitted to motion correction using MCFLIRT (Jenkinson et al. 2002). Structured artefacts were removed by combining ICA and FMRIB’s ICA-based X-noiseifier (Griffanti et al., 2014). To help reduce unnecessary interpolation effects, all intermediate warp operations were merged into a composite transformation allowing for simultaneous application to fMRI maps. For the display of results (see Figures 1-3), maps were projected to the cortical surface. This was done via volume-to-surface mapping in wb_command (www.humanconnectome.org), based on the Human Connectome Project (HCP) group average template “S1200_MSMAll”.

**Figure 1.**
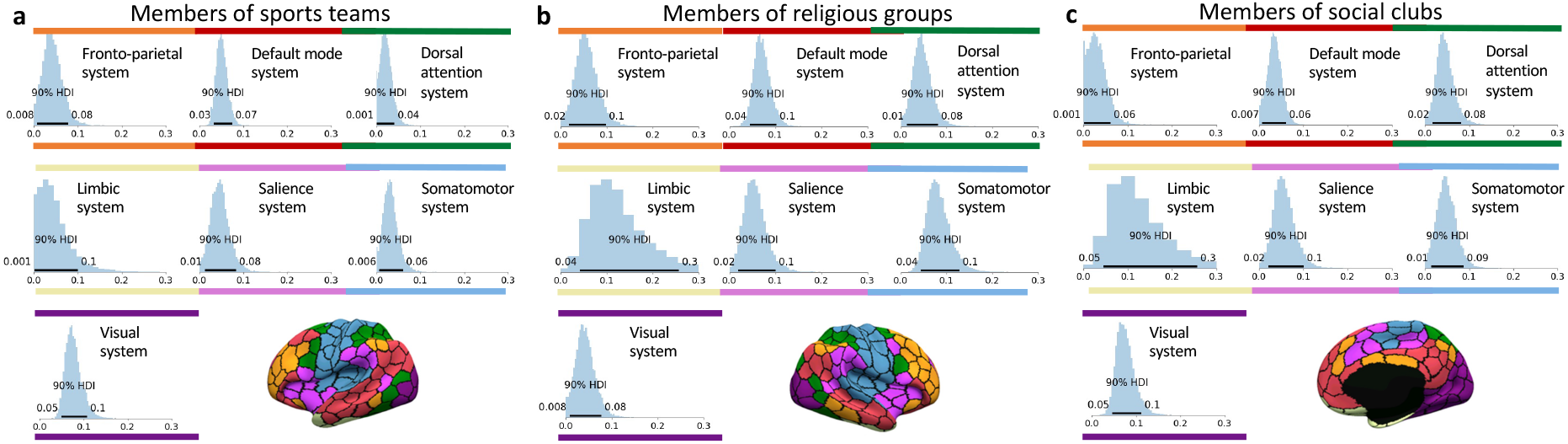
Three forms of social participation show strong network-wide effects in the default mode system and limbic systems. A Bayesian hierarchical framework was used to jointly analyze overall volume variation in seven major brain networks at the population-level (UK Biobank cohort with n = ~ 40.000 participants), separately to identify members of sports teams (**a**), religious groups (**b**) and social clubs (**c**). Contributions exhibit the amount of variance explained by every distributed canonical network, according to the 10-90% highest posterior density interval (HDI). A narrow (wide) posterior parameter distribution posterior density stands for a certain (uncertain) effect. A more positive (less positive) value for the mean posterior parameter density (x axis, sigma parameter) indicates for a higher (lower) explained variance across the dependent spatially distributed atlas regions in that network. By rough analogy to classical ANOVA, the network definitions could be viewed as “factors” and the region definitions could be viewed as “levels”. Results are adjusted for effects of age and sex.

**Figure 2.**
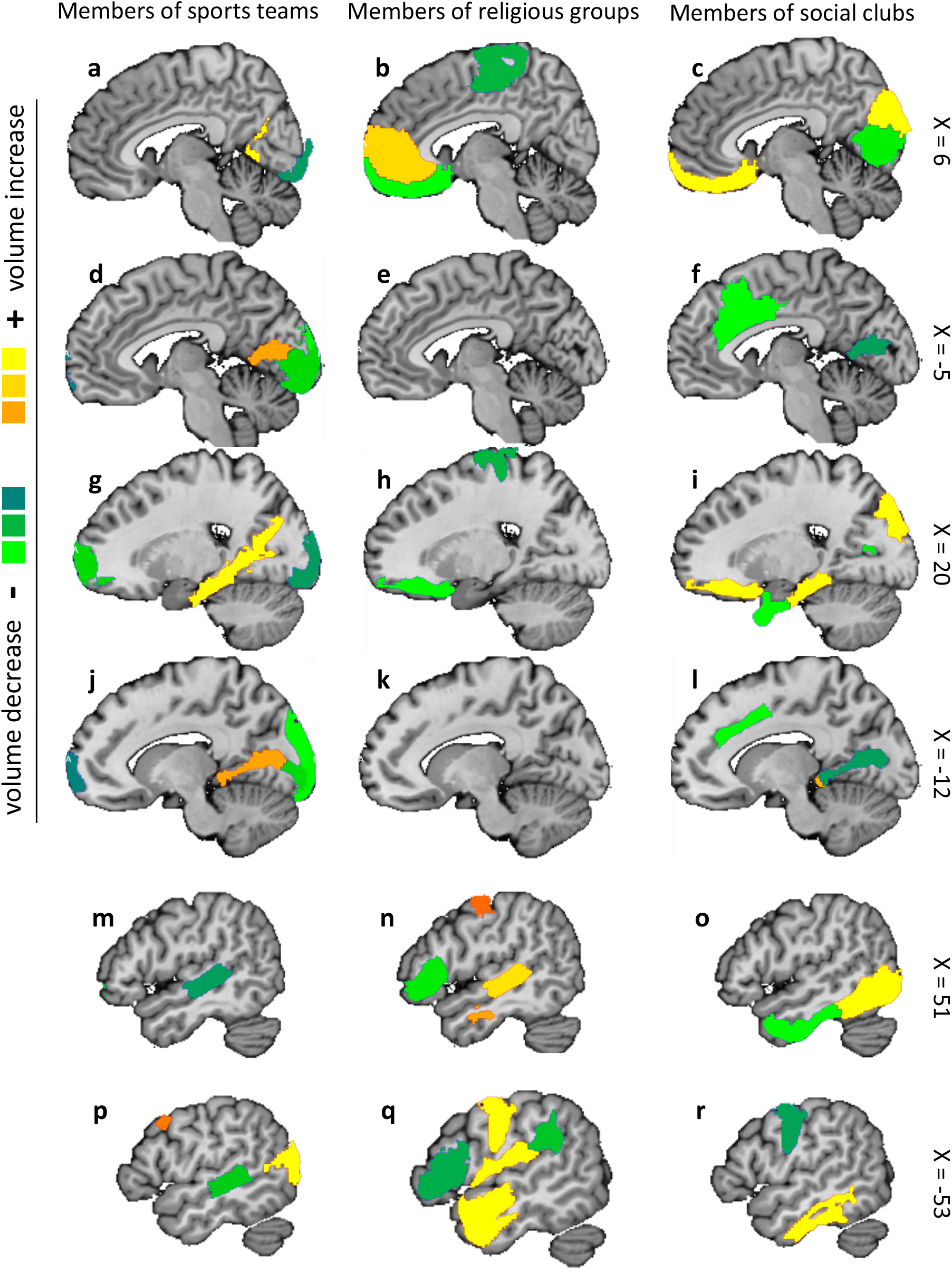
Population associations with regions of the default and limbic systems dominate the neural correlates of social participation. At the region level, our Bayesian hierarchical model identified for which brain regions variability in grey matter volume explains participants’ reported weekly engagement in social groups. Strongest associations to day-to-day social support (cf. Fig. 1) were determined based on effect sizes (mean parameter) of the marginal posterior parameter distributions (volume measures in standard units). In the context of our modeling solution, the ventromedial prefrontal cortex showed no volume deviation in right hemisphere of (**a**) sport teams participants. A ventromedial prefrontal gray matter decrease was shown in (**b**) religious groups (light green) and a substance increase in participants of (**c**) social clubs (yellow). The anterior cingulate cortex showed no volume deviation in (**d**) sports team members in members of (**e**) religious groups in the left hemisphere. A volume decrease in the anterior cingulate cortex was instead shown in members of (**f**) social clubs (light green). The parahippocampal area showed a volume increase within the group of (**g**) sports teams (yellow) and (**i**) social clubs (yellow), while no volume deviation was shown for (**h**) religious groups. For the lingual gyrus, positive and negative volume effects were shown for (**j**) sports teams, and no volume effects for (**k**) religious groups. Negative volume effects were shown for (**l**) social clubs. Within the temporal lobe of sports team members (**m, p**), both hemispheres had negative volume effects in the superior and middle temporal gyrus (green). For members of religious groups (**n, q**), volume decrease in both hemispheres was demonstrated in superior and middle temporal gyrus (yellow). The social club participants (**o, r**) showed a volume decrease (green) in right hemisphere and volume increase in left hemisphere in fusiform gyrus and inferior temporal gyrus.

**Figure 3.**
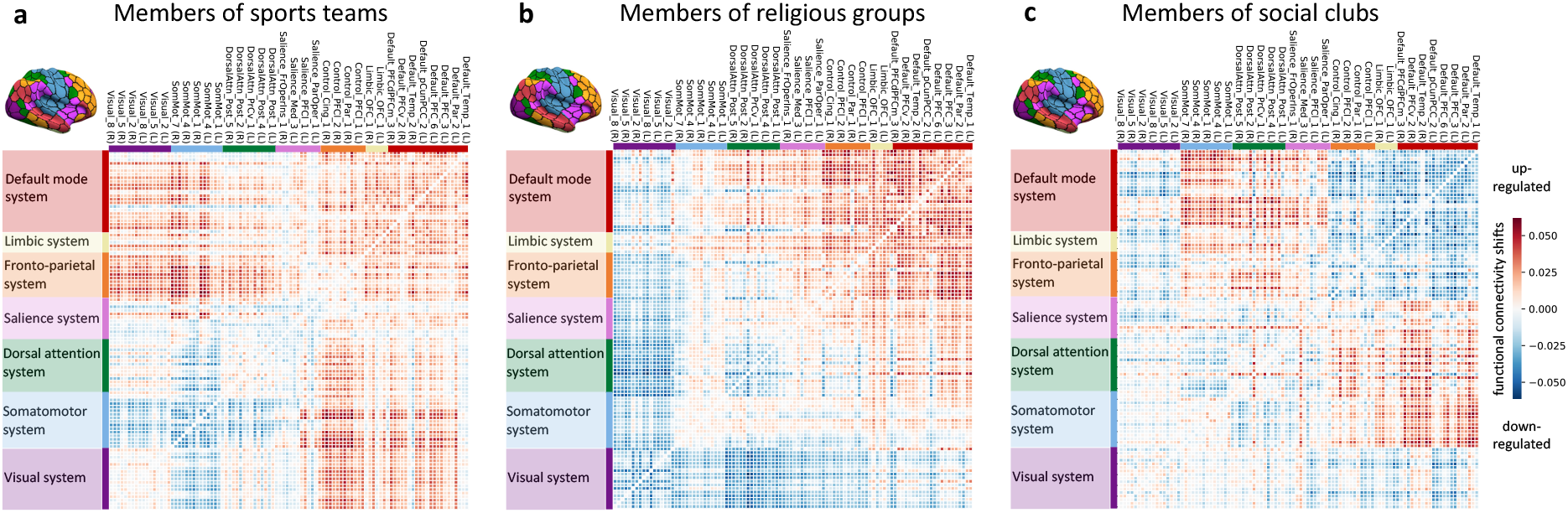
Signatures of functional coupling fluctuations highlight the relevance of intra- and inter-network deviations of the default and limbic network in social participation. Functional connectivity shifts are shown for the dominant population mode related to weekly social participation (connectivity links in standard units). The leading mode identified positive (red) and negative (blue) shifts in network connectivity using a pattern-learning algorithm, with statistical significance at *p* < 0.05 (one-sided test) using nonparametric permutation testing. The intra- and inter-network connectivity is depicted for the participants of (**a**) sports teams, (**b**) religious groups and (**c**) social clubs. L/R refers to left and right hemisphere. The default and limbic network showed an increase in connectivity strengths in members of sport teams and religious groups and a decrease in members of social clubs.

### Analysis of associations between social participation and grey matter patterns

Neurobiologically interpretable measures of grey matter volume were extracted in all participants by summarizing whole-brain sMRI maps in Montreal Neurological Institute (MNI) reference space. This feature generation step was guided by the topographical brain region definitions of the widely used Schaefer-Yeo atlas comprising 100 parcels (Schaefer et al. 2018). The participant-level brain region volumes provided the input variables for our Bayesian hierarchical modeling approach (cf. below). As a data-cleaning step, inter-individual variation in brain region volumes that could be explained by variables of no interest were regressed out: body mass index, head size, average head motion during task-related brain scans, average head motion during task-unrelated brain scans, head position and receiver coil in the scanner (x, y, and z), position of scanner table, as well as the data acquisition site.

To examine population variation of our atlas regions in the context of regular social participation, we have purpose-designed a Bayesian hierarchical model, building on our previous research (Bzdok et al., 2017; Bzdok et al., 2020; Kiesow et al., 2020). In contrast, classical linear regression combined with statistical significance testing would simply have provided p-values against the null hypothesis of no difference between individuals with high and low social participation in each brain region. Instead of limiting our results and conclusions to strict categorical statements, each region being either relevant for differences in social participation or not, our analytical strategy aimed at full probability distributions that expose how brain region volumes converge or diverge in their relation to social participation as evidenced in the UK Biobank population. In a mathematically rigorous way, our approach estimated coherent, continuous estimates of uncertainty for each model parameter at play for its relevance in social participation. Our study thus addressed the question “How certain are we that a regional brain volume is divergent between high and low social participation individuals?”. Our analysis did not ask “Is there a strict categorical difference in region volume between high and low social participation individuals?”.

The elected Bayesian hierarchical framework also enabled simultaneous modeling of multiple organizational principles: i) *segregation* into separate brain regions and ii) *integration* of groups of brain regions in form of spatially distributed brain networks. Two regions of the same atlas network are more likely to exhibit similar volume effects than two regions belonging to two separate brain networks. Each of the region definitions was pre-assigned to one of the 7 large-scale network definitions in the Schaefer-Yeo atlas (Schaefer et al. 2018) or the collection of subcortical regions from the Harvard-Oxford atlas (Desikan et al., 2006), providing a native multilevel structure. Setting up a hierarchical generative process enabled our analytical approach to borrow statistical strength between model parameters at the higher network level and model parameters at the lower level of constituent brain regions. By virtue of exploiting partial pooling, the brain region parameters were modeled themselves by the hyper-parameters of the hierarchical regression as a function of the network hierarchy to explain social participation. Assigning informative priors centered around zero provided an additional form of regularization by shrinking coefficients to zero in the absence of evidence to the contrary. We could thus provide fully probabilistic answers to questions about the morphological relevance of individual brain locations and distributed cortical networks by a joint varying-effects estimation that profited from several biologically meaningful sources of population variation.

The model specification placed emphasis on careful inference of unique posterior distributions of parameters at the brain network level to discriminate individuals with (encoded as outcome 0) and without (outcome 1) a certain social group membership:

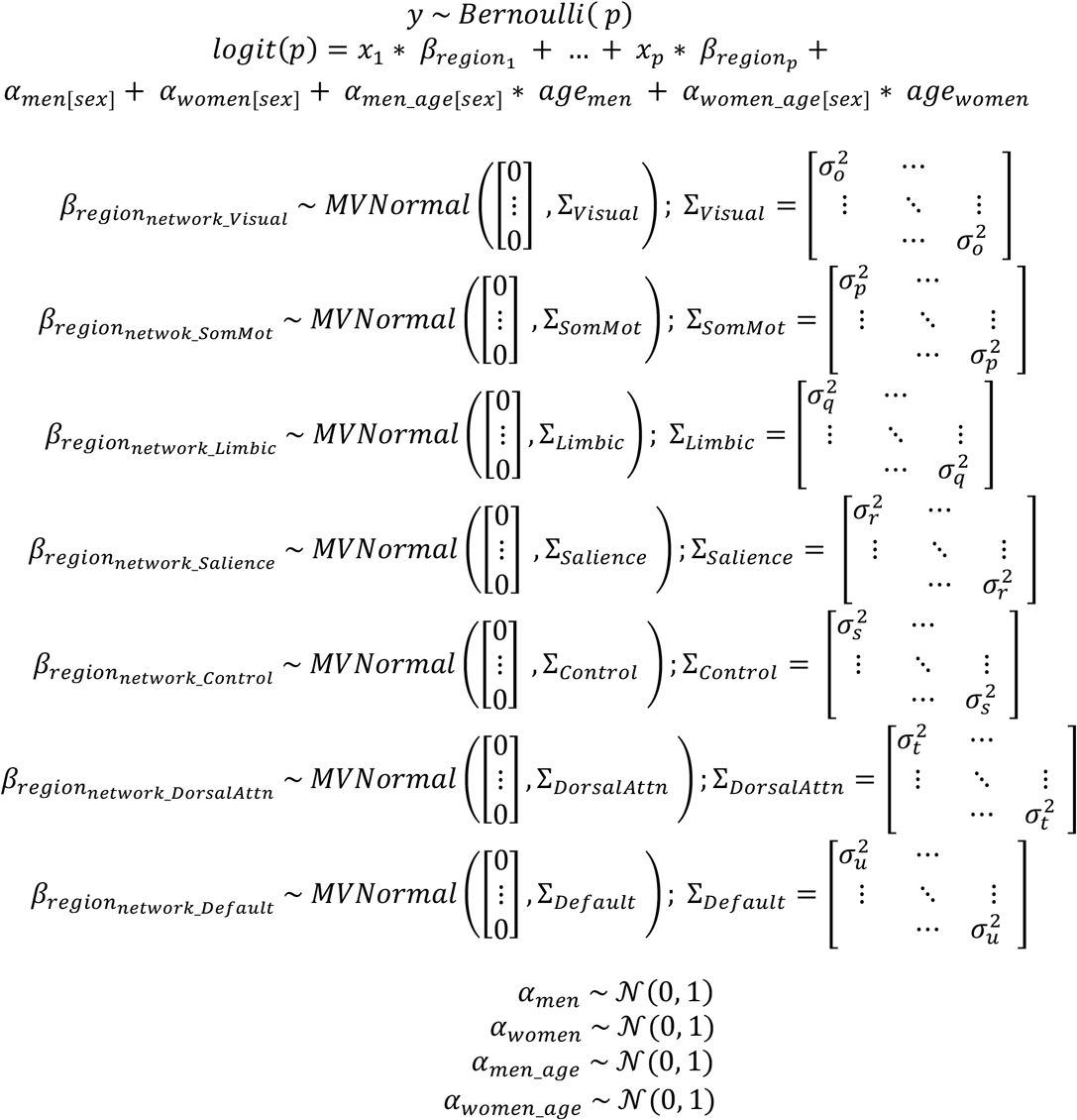

where *sigma* parameters estimated the overall variance across the *p* brain regions that belong to a given atlas network, independent of whether the volume effects of the respective constituent brain regions had positive or negative direction. As such, the network variance parameters *sigma* directly quantified the magnitude of intra-network coefficients, and thus the overall relevance of a given network in explaining regular social participation based on the dependent region morphology measures. All regions belonging to the same brain network shared the same variance parameter in the diagonal of the covariance matrix, while off-diagonal covariance relationships were zero.

Probabilistic posterior distributions for all model parameters were estimated for the hierarchical models. Our Bayesian approach could thus simultaneously appreciate grey matter variation in segregated brain regions as well as in integrative brain networks in a population cohort. The approximation of the posterior distributions was carried out by the NUTS sampler (Gelman et al., 2014), a type of Markov chain Monte Carlo (MCMC), using the PyMC3 software suite (Salvatier et al., 2016). After tuning the sampler for 4,000 steps, we drew 1,000 samples from the joint posterior distribution over the full set of parameters in the model for analysis. Proper convergence was assessed by ensuring Rhat measures (Gelman et al. 2014) stayed below 1.02.

### Analysis of associations between social participation and functional connectivity patterns

Quantitative measures of functional connectivity were computed for cortex-wide brain regions as defined by the Schaefer-Yeo atlas (Schaefer et al. 2018). Functional connectivity profiles for each participant were derived by computing Pearson’s correlation between their neural activity fluctuations. To this end, in each participant, the time series of whole-brain fMRI signals, obtained in the absence of an externally structured experimental task, were summarized by averaging for each brain region in the atlas. The approach yielded the functional coupling signature of the whole cortex as a 100 x 100 region coupling matrix for each participant. The ensuing region-region coupling estimates underwent standardization across participants by centering to zero mean and unit scaling to a variance of one (cf. next step). Inter-individual variation in the functional coupling strengths between brain regions that could be explained by variables of no interest were regressed out in a data-cleaning step (analogous to sMRI analysis): body mass index, head size, average head motion during task-related brain scans, average head motion during task-unrelated brain scans, head position as well as receiver coil in the scanner (x, y, and z), position of scanner table, and data acquisition site, as well as age, sex and age-sex interactions.

We then sought the dominant coupling regime – signature or “mode” of population covariation – that provides insight into how functional variability in 100 brain regions can explain regular social participation. Partial least squares (PLS) was an ideal analytical method to decompose the obtained 100 x 100 fingerprint matrix of functional couplings with respect to social participation. The variable set *X* was constructed from the lower triangle of the participants’ functional coupling matrices. The target vector *y* encoded more socially engaged participants as +1 and participants without a given social group membership as −1. PLS involves finding the matrix factorization into *k* low-rank brain representations that maximize the correspondence with our social trait of interest. PLS thus identified the matrix projection that offered the maximal covariance between sets of region couplings in the context of participant reports of social participation.

In other words, the extracted functional coupling mode identified the driving linear combinations of cortical brain connections that featured the best correspondence to regular social participation. Concretely, positive (negative) modulation weights revealed increased (decreased) correlation strengths, relative to average functional coupling. This is because the computed functional connectivity estimates were initially normalized to zero mean and unit variance across participants. For example, a functional connectivity input into PLS of 0 denoted the average functional coupling strength in our UK Biobank sample, rather than an absence of functional connectivity between the region pair. The derived pattern of PLS weights, or canonical vectors, thus indicated deviations from average functional coupling variation in our cohort. Moreover, the variable sets were entered into PLS after a confound-removal procedure (cf. above).

Next, we assessed the statistical robustness of the resulting dominant PLS mode of functional coupling deviations related to social participation in a non-parametric permutation procedure, following previous research (Miller et al. 2016). Relying on minimal modeling assumptions, a valid empirical null distribution was derived for the Pearson’s correlation between low-rank projections of the dominant mode resulting from PLS analysis. In 1,000 permutation iterations, the functional connectivity matrix was held constant, while the social participation labels were submitted to random shuffling. The constructed surrogate datasets preserved the statistical structure idiosyncratic to the fMRI signals, yet permitted to selectively destroy the signal properties that are related to social participation (Efron, 2012). The generated distribution of the test statistic reflected the null hypothesis of random association between the brain’s functional coupling and regular social participation across participants. We recorded the Pearson’s correlations rho between the perturbed low-rank projections in each iteration. P-value computation was based on the 1,000 Pearson’s rho estimates from the null PLS model.

### Demographic profiling analysis of the brain correlates of social participation

We finally performed a profiling analysis of the brain regions that were most strongly associated with regular social participation. Based on our results in brain structure, we carried out a rigorous test for multivariate associations between our top regions and a diverse set of indicators that exemplify the domains of a) basic demographics, b) personality features, c) substance-use behaviours, and d) social network properties (for details see https://www.ukbiobank.ac.uk/data-showcase/). The set of behavioural variables and the set of brain measures were z-scored across participants to conform to zero mean and unit variance. The brain variables were submitted to the top 10 (sMRI) of brain measures that were identified as most important in the context of social participation (cf. above). In the case of brain structure, the target brain regions were selected based on the (absolute) modes of the Bayesian posteriors of marginal parameter distributions at the region level (cf. above). In the case of brain function, the target brain connections were selected based the (absolute) effect sizes from the dominant PLS mode (cf. above).

Using the two variable sets of brain and behaviour measurements, we then carried out a bootstrap difference analysis of the collection of target traits in individuals with high versus low social participation (Efron & Tibshirani, 1994). In 1,000 bootstrap iterations, we randomly pulled equally sized participant samples to perform a canonical correlation analysis (CCA), in parallel, in individuals with and without social group membership (Miller et al., 2016; Wang et al., 2020). In each resampling iteration, this approach estimated the doubly multivariate correspondence between the brain and behaviour indicators in each group. The ensuing canonical vectors of the dominant CCA mode indicated the most explanatory demographic associations in a given pull of participants. To directly estimate resample-to-resample effects in group differences, the canonical vectors of behavioural rankings were subtracted elementwise between participants from both groups, recorded, and ultimately aggregated across the 1,000 bootstrap datasets.

This analytical tactic allowed propagating the noise of participant sampling variation into the computed uncertainty estimates of group differences in the UK Biobank cohort. Statistically defensible behavioural dimensions were determined by whether the (two-sided) bootstrap confidence interval included zero or not in the 5/95% bootstrap population interval. In a fully multivariate setting, this non-parametric modeling scheme directly quantified the statistical uncertainty of how a UK Biobank trait is differentially linked to brain-behaviour correspondence as a function of regular social participation.

## Results

We have mined the UK Biobank resource with a focus on social participation that indicate weekly attendance in a sports team, religious group or social club. Brain region information on structural volume and functional coupling was extracted in ~40,000 UK Biobank Imaging participants guided by the Schaefer-Yeo reference atlas (Schaefer et al., 2018). All results have adjusted the brain features for variation that could be explained by age and sex/gender (cf. methods).

### Structural brain correlates of sports team participation

At the network level, in participants who attend sports teams at least once every week, the highest explanatory relevance across spatially distributed brain regions emerged in the visual network (posterior sigma = 0.08; 10-90% highest posterior density [HPD] = 0.05/0.10), followed by the DMN (sigma = 0.05, HPD = 0.04/0.07) and the limbic network (sigma = 0.05, HPD = 0.01/0.08). Further volume associations for sports team members were located to (in descending order, based on the explanatory quality): the salience network (sigma = 0.05, HPD = 0.02/0.07), frontoparietal control network (sigma = 0.05, HPD = 0.01/0.07), somatomotor network (sigma = 0.03, HPD = 0.01/0.05) and dorsal attention network (sigma = 0.02, HPD = 0.01/0.04). Overall, the largest but most uncertain brain-behaviour association was featured by the limbic network. In contrast, as indicated by the narrowness of the inferred network-level posterior parameter distribution, the network with the most certain contribution to explaining volume variation in the context of sports team membership was the DMN (Fig. 1).

At the region level of the obtained Bayesian hierarchical modeling solution, positive volume effects emerged especially in the left lingual gyrus (posterior mean = 0.05, 10-90% HPD = 0.01/0.10), right posterior cingulate cortex (mean = 0.08, HPD = 0.03/0.14), left middle temporal gyrus (mean = 0.09, HPD = 0.04/0.14), left middle frontal gyrus (LH_Default_PFC_6; mean = 0.05, HPD = 0.01/0.08 and LH_Default_PFC_7; mean = 0.07, HPD = 0.03/0.11) and right parahippocampal gyrus (mean = 0.09, HPD = 0.03/0.16). Negative associations between volume of brain regions and sports team membership were found for the left (mean = −0.10, 10-90% HPD = −0.14/-0.05) and right inferior occipital gyrus (mean = −0.06, HPD = −0.11/-0.01), left lingual gyrus (mean = −0.09, HPD = −0.13/-0.04), left middle occipital gyrus (mean = −0.10, HPD = −0.16/-0.05), left middle temporal gyrus (mean = −0.08, HPD = −0.13/0.03), left (mean = −0.05, HPD = −0.10/-0.01) and right superior frontal gyrus (mean = −0.09, HPD = −0.14/-0.01), as well as right superior temporal gyrus (mean = −0.06, HPD = −0.11/-0.01). Overall, we found regions of the default and limbic network as well as lingual gyrus, prefrontal and temporal cortex to be the key brain correlates linked to weekly engagement in a sports team.

### Structural brain correlations of religious group participation

At the network level, for people who attend religious groups on a weekly basis, we found the strongest association between regular social participation and volume variation for the limbic network (posterior sigma = 0.15, 10-90% HPD = 0.05/0.21), somatomotor network (sigma = 0.09, HPD = 0.05/0.11) and the DMN (sigma = 0.07, HPD = 0.05/0.09). In descending order of explanatory quality, the frontoparietal control network (sigma = 0.06, HPD = 0.03/0.09) and salience network (sigma = 0.06, HPD = 0.03/0.09) showed relevant effects, accompanied by dorsal attention network (sigma = 0.05, HPD = 0.02/0.07) and visual network (sigma = 0.04, HPD = 0.02/0.07). Similar to our hierarchical pattern analysis on sports teams (cf. above), the prominent effect with greatest uncertainty was seen in the limbic network, while the most certain network-level volume effect was located to the DMN.

Regarding region-level effects of the obtained Bayesian hierarchical modeling solution, we identified several relevant regions that characterize regular attendees of religious groups. Positive volume effects became apparent in the left postcentral gyrus (posterior mean = 0.10, 10-90% HPD = 0.02/0.17), left precentral gyrus (LH_SomMot_4; mean = 0.14, HPD = 0.07/0.20; LH_Default_Temp_1; mean = 0.11, HPD = 0.03/0.18), right precentral gyrus (mean = 0.06, HPD = 0.01/0.12), right middle temporal gyrus (mean = 0.08, HPD = 0.01/0.14), right superior temporal gyrus (mean = 0.10, HPD = 0.03/0.16) and right rostral anterior cingulate cortex (mean = 0.09, HPD = 0.02/0.16). Furthermore, we identified a set of regions that showed a negative association with religious group participation: left inferior parietal lobule (mean = −0.08, HPD = −0.13/-0.01), left inferior frontal gyrus (mean = −0.07, HPD = −0.13/-0.01), right paracentral lobule (mean = −0.07, HPD = −0.13/-0.01), right dorsal anterior cingulate cortex (mean = −0.16, HPD = −0.26/-0.06) and right inferior frontal gyrus (mean = −0.09, HPD = −0.15/-0.03). As such, similar to our results for sports team members (cf. above), the default and limbic network were highlighted in spiritually active people.

### Structural brain correlates of social club participation

At the network level, social participation in the form of weekly attendance of social clubs was also linked to a collection of grey matter deviations of atlas region sets. In this third and last analysis of social participation, the largest explained variance and also the most uncertain effect was found for the limbic network (posterior sigma = 0.15, 10-90% HPD = 0.05/0.20). The following networks were less explanatory (in descending order): the visual network (sigma = 0.08, HPD = 0.05/0.10), salience network (sigma = 0.06, HPD = 0.03/0.09), somatomotor network (sigma = 0.05, HPD = 0.02/0.08), dorsal attention network (sigma = 0.05, HPD = 0.02/0.07), DMN (sigma = 0.04, HPD = 0.01/0.06) and control network (sigma = 0.03, HPD = 0.01/0.05). The most informative distribution among the seven networks for social club attendance is the visual network, followed closely by the default network.

At the region-level of the obtained Bayesian hierarchical modeling solution, we found volume effects characterizing social club members for the atlas region sets. A positive effect for grey matter volume was found for the left (posterior mean = 0.07, 10-90% HPD = 0.01/0.13) and right parahippocampal gyrus (mean = 0.11, HPD = 0.04/0.19), left inferior temporal gyrus (mean = 0.11, HPD = 0.05/0.17), right fusiform gyrus (mean = 0.11, HPD = 0.04/0.17), right cuneus (mean = 0.09, HPD = 0.03/0.14) and right anterior cingulate cortex (mean = 0.11, HPD = 0.03/0.19). We identified the most relevant negative volume deviations for social club participants in the left lingual gyrus (mean = −0.06, HPD = −0.11/-0.01), left precentral gyrus (mean = −0.06, HPD = −0.11-/0.01), left dorsal and ventral anterior cingulate cortex (mean = −0.11, HPD = −0.18-/0.04), right lingual gyrus (mean = −0.10, HPD = −0.15/-0.03) and right middle temporal gyrus (mean = −0.13, HPD = −0.22/-0.04). In agreement with our findings for sports team and religious group members, regions of the limbic and default network were again found to be prominent (Table 1). In addition, among all three groups, similar regions tended to come to the fore, including parahippocampal and fusiform gyrus, anterior cingulate cortex, temporal and prefrontal cortex (Fig. 2).

**Table 1.**
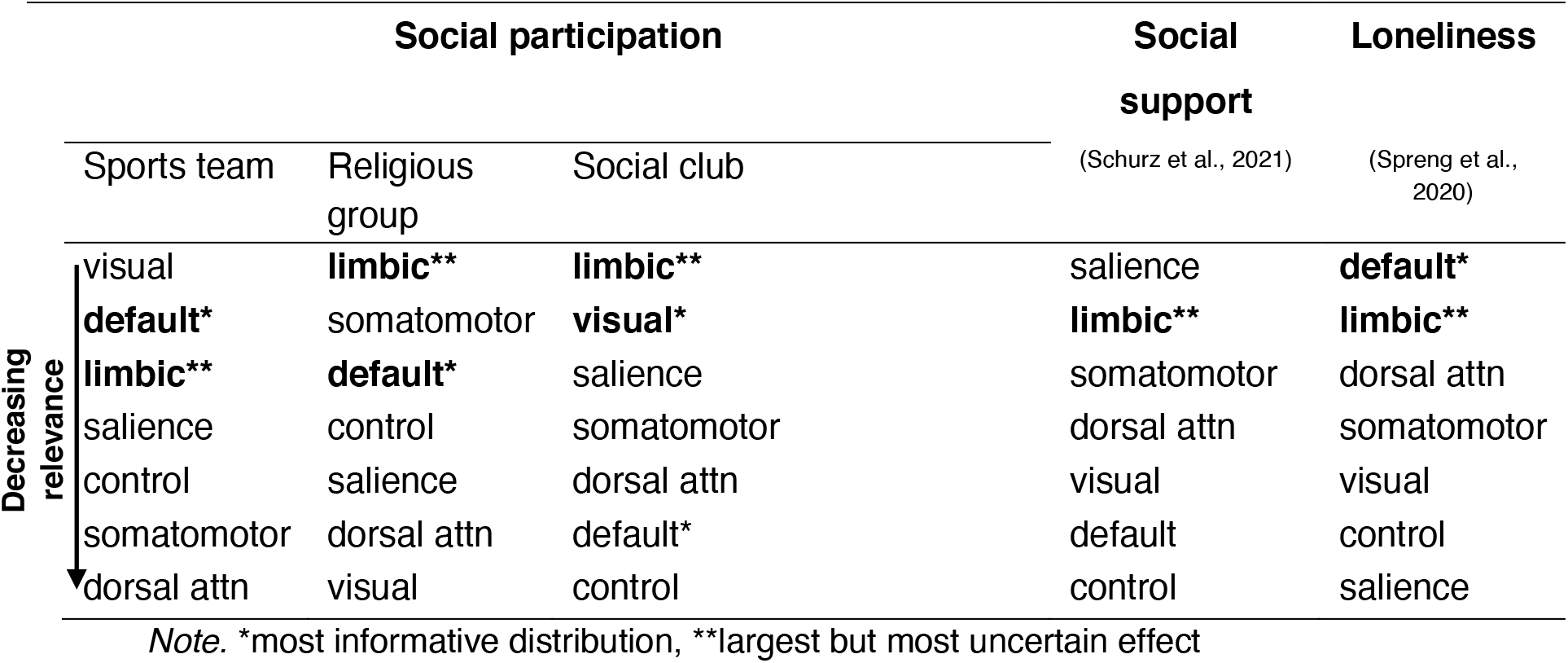
List of analyzed 7 networks, based on Schaefer-Yeo atlas, in decreasing order of explained variance (top to bottom) based on highest posterior density (HPD).

### Functional brain correlates of social participation

Next, fMRI data from our UK Biobank cohort were examined to investigate possible deviations in functional coupling fingerprints related to weekly engagement in sports teams, religious groups and social clubs. The cortex-wide functional connectivity profiles of each participant were submitted to a multivariate pattern-learning algorithm that identified a collection of reliable positive and negative shifts in network connectivity in the context of social participation (*p* < 0.05). In so doing, the single most coherent pattern of deviation for the functional connectome of participants within each of the three groups was identified (Fig. 3).

The regular attendees of sports teams showed wide-ranging deviations in intra-network connectivity of default and limbic network. The somatomotor network revealed negative coupling effects in intra-network connectivity. An increase in inter-network connectivity was dominated by connections from the DMN, but also featured the limbic and frontoparietal control network’s functional ties to the somatomotor network. Strengthened coupling was detected between the DMN and most other examined large-scale functional networks, including visual network, somatomotor network, frontoparietal control network and limbic network. The frontoparietal control network was found to exhibit enhanced coupling links especially with both the visual and somatomotor networks. A decrease in functional coupling strength was observed for the somatomotor network, the visual network and the dorsal attention network.

People participating in religious groups in turn were especially characterized by a compounding of within-network functional connections within the DMN, limbic network and to some extent also in the frontoparietal control network. These three neural network systems showed enhanced within- and between-network functional connectivity patterns. In contrast, the dorsal attention network and the visual network both showed reduced within-network connectivity strengths. Connectivity strengths between regions of the DMN and the neural circuits of the limbic, frontoparietal control and dorsal attention network were significantly increased. Furthermore, we identified a decrease in connectivity strengths between the visual network and several other large-scale networks, including dorsal attention network, frontoparietal control network, salience network as well as the somatomotor network to a reduced extent.

In contrast to these two types of social participation, social participation in social clubs did not lead to a salient increase of functional connectivity strengths within most of the aforementioned networks. Instead, a relevant decrease in functional connectivity was noted within DMN and limbic networks as the single most coherent pattern of deviation. The DMN in turn showed enhanced functional coupling with the somatomotor and the dorsal attention networks.

As such, among all three types of social participation, the default and the limbic network stuck out in the overall collection of observed deviations in region-region coupling strengths. This insight was evidenced by enhanced connectivity patterns in active members of sports teams and religious groups, and by diminished connectivity patterns in social club participants. Additionally, reminiscent of our general findings from the structural analyses (cf. above), the default and limbic network played a prominent role among the systematic shifts of between-network coupling.

### Demographic profiling analyses of social participation

In the final set of analyses, we have linked grey matter volume deviation in the most relevant identified regions (top 10%) to behavioural and sociodemographic data via a multivariate pattern analysis in each of the three groups of social participation (Fig. 4). In terms of consistent findings across groups, the time spent watching television ranked highest in all three separate analyses: sports teams (mean = −0.39, 5/95% confidence interval [CI] = −0.90/-0.07), religious groups (mean = −0.44, CI = −0.73/-0.08) and social clubs (mean = −0.56, CI = −0.93/-0.16). Another of our findings that were broadly consistent across all three groups pertained to the number of persons in the closest family circle: the number of siblings, brothers and sisters, showed high concordance across members of sports teams (mean_sisters_ = −0.20, 5/95% CI = −0.66/0.10; mean_brothers_ = −0.16, CI = −0.71/0.15), of religious groups (mean_sisters_ = −0.40, CI = −0.73/0.01; mean_brothers_ = −0.41, CI = −0.77/-0.01) and of social clubs (mean_sisters_ = −0.32, CI = −0.68/0.06; mean_brothers_ = −0.31, CI = −0.66/0.06). Group membership was also reflected by similarities in health-related lifestyle behaviours. Again, all three forms of social participation showed similar convergence for the type and extent of the consumption of alcohol intake (amount of alcohol drunk on a typical drinking day, alcohol intake frequency) and tobacco use (past tobacco smoking, current tobacco smoking). To a varying extent, different psychological conditions also showed a high concordance within all three groups of social participation. These psychological conditions included including loneliness, mood swings, neuroticism, fed-up feelings, suffer from “nerves”, tense, miserableness, irritability and sensitivity. In sum, charting relevant brain-behaviour associations revealed a high concordance among all three forms of social participation on sociodemographic and behavioural level for family structure, alcohol and tobacco consumption.

**Figure 4.**
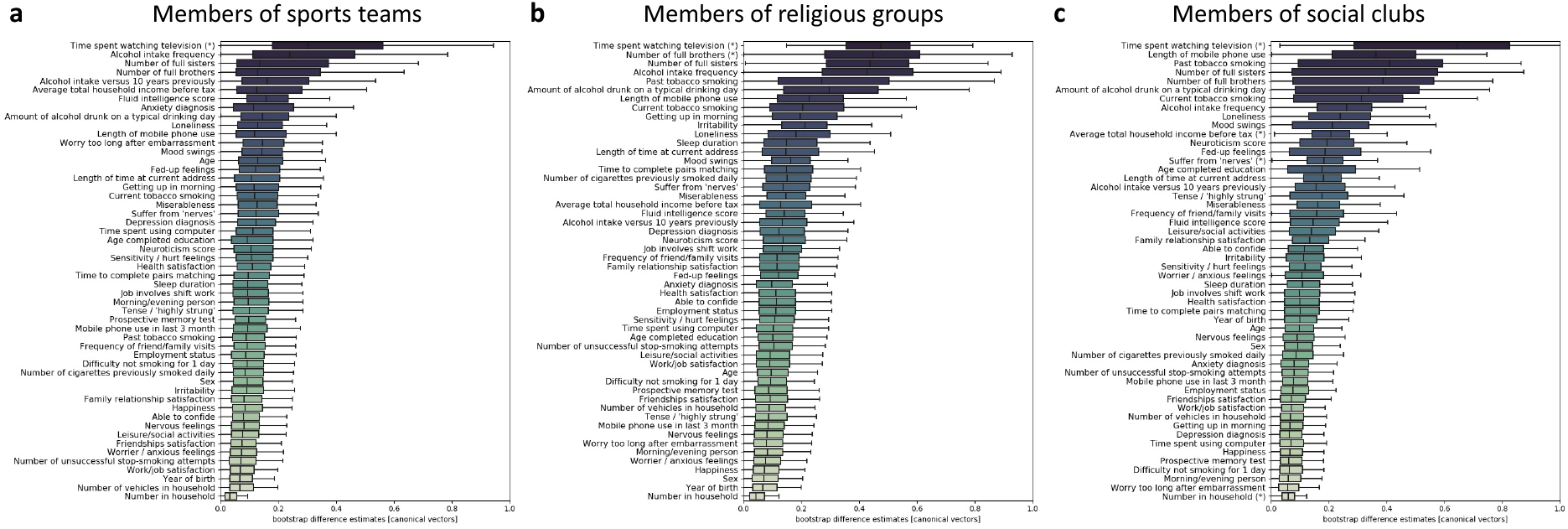
Demographic profiling analysis pinpoints indicators of mental health, smoking and alcohol consumption consistently across three groups of social participation. Multivariate pattern-learning (cf. Methods) was used to explore how the top brain regions (see Fig. 2) are linked to a variety of behavioral indicators with their robust cross-links to weekly social participation. Behavioral measures covered domains of mental and physical well-being, lifestyle choices, and social embeddedness. In 1,000 bootstrap resampling iterations, our entire pattern-learning pipeline in grey matter volume was repeated separately in the two participant groups: UK Biobank participants who regularly share life experience with close others and those with little such exchange of personal events. The computed differences in brain-behavior associations between both groups (i.e., dominant canonical vector entries) were gathered across the 1,000 perturbed realizations of our original dataset to obtain faithful bootstrap intervals. These estimates of uncertainty directly quantified how group-related deviations vary in the wider population. Asterisks indicate statistical relevance based on excluding zero between the 5/95% quantiles of the bootstrap distribution. These brain-behavior associations showed great correspondence for regular intake of alcohol and tobacco as well as multifaceted aspects of psychological well-being across participants of (**a**) sports teams, (**b**) religious groups and (**c**) social clubs. Further consistent findings across all three forms of social participants included the time spent watching television, number of siblings and further mental health conditions. The boxplot whiskers depict the interquartile range.

## Discussion

Experiencing times of unmet social desire and not being able to fulfil one’s need to belong can be a wake-up call and highlights the pivotal importance of community and social exchange. Consequently, exploring the neurobiological substrates of social participation and their ties to physical and mental health is imperative. In our present investigation, the DMN and the limbic system were placed in the center of robust brain manifestations across three examined types of social participation in ~40,000 UK Biobank participants: sports teams, religious groups, and social clubs. First, our structural analyses demonstrated the importance of prefrontal and cingulate cortex. Parts of the ventromedial prefrontal cortex (vmPFC) emerged as particularly relevant for participation in all three social groups. Second, in our analyses of functional coupling fingerprints, the DMN and limbic system also emerged consistently among all three groups as the cornerstone of the neural bases indicative of social embeddedness via group membership. Third, the combination of neuroimaging with behavioural and sociodemographic data showed a high consistency among all three types of participation for alcohol and tobacco consumption as well as certain psychological states. Hence, participants of sport teams, religious groups and social clubs do not only show similarities on the neural level, but also on the level of substance use behaviour and mood.

The close involvement of the DMN in social and affective processes has been reported on multiple occasions (Amft et al., 2015; Amodio & Frith, 2006; Hyatt et al., 2015; Mars et al., 2012; Redcay & Schilbach, 2019; Schilbach et al., 2008). The relevance of the DMN in subserving advanced social capacities in various contexts has repeatedly been highlighted based on studies of both functional connectivity and grey matter volume (Che et al., 2014; Chen et al., 2010; Finlayson-Short et al., 2020; Lee, 2014; Morita et al., 2021; Takeuchi et al., 2014). Volume variation in the DMN related to social cognition has been reported for the prefrontal cortex as well as the anterior cingulate cortex (Morita et al., 2021) and posterior cingulate cortex (Che et al., 2014). Volume associations with the limbic system were previously linked to the extent of social support and satisfying friendship bonds (Taebi et al., 2020). Extending these previous findings, our present study reveals volume deviation and shifts in functional connectivity patterns that center on the DMN. We linked this neural system to real-world membership in a sports team, religious group or social club on the population level for the first time. This broadens the interpretational perspective on the DMN by adding relevant aspects of everyday life that cannot usually be studied in experimental MRI research.

Looking at specific regions related to the DMN, we found that the vmPFC showed a positive volume effect in participants of social clubs and a negative volume effect in participants of sports teams and religious groups. Lesion studies as well as neuroimaging studies in humans and animals pointed out the important role of the vmPFC in complex forms of social cognition, such as mentalization, empathy and decision making (Hiser & Koenigs, 2018; Leopold et al., 2012; Shamay-Tsoory et al., 2003). Larger volume of the vmPFC has been directly linked to the size of an individual’s social network and mentalizing competences (Lewis et al., 2011; Powell et al., 2012).

The anterior cingulate cortex (ACC) and parahippocampal gyrus are often attributed to the limbic system. The parahippocampal gyrus revealed additional grey matter volume within sports team and social club participants. The ACC showed significant volume deviations for religious groups and social clubs but none for sport teams. For religious groups, ventral parts of ACC showed positive volume effects, while dorsal parts showed negative volume effects. For social club members, dorsal and ventral volume effects for cingulate cortex differed between the two hemispheres. Given that social behaviour is potentially most uniquely developed in the human species (Frith & Frith, 2010) it may come as no surprise to find brain asymmetry features, as hemispheric asymmetry is also exceptionally well developed in the human brain (Hartwigsen et al., 2021). The regional findings match the notable interindividual variability of the limbic-trait associations at the network-level of our Bayesian hierarchical analyses. Only recently, grey matter volume increase in the dorsal and perigenual ACC was linked to *social affective benefit* (Gan et al., 2021). The community-based cohort study recorded daily-life social contacts and affective valence via smartphone for one week and combined the data with neuroimaging. Applied to the current results, this might indicate less social affective benefit from the membership in religious groups.

The fusiform gyrus is classically associated with processing information from other people’s faces. These neural processes assist in recognizing a person identity from their unique facial features – hence, especially stable features of other’s faces (Schilbach et al., 2012). The posterior superior temporal sulcus in turn is repeatedly linked to basic sensory input relevant for processing social interactions (right hemisphere; Isik et al., 2017; Walbrin et al., 2018) and also to face processing (Bzdok et al., 2011; Bzdok et al., 2012). Furthermore, this region is classically linked to processing variable facial features like eye gaze and emotional cues.

Previous studies compared the neuronal activation of fusiform gyrus and posterior superior temporal sulcus through trustworthiness, attractiveness, emotion and age judgments (Bzdok et al., 2012; Oosterhof & Todorov, 2008). While posterior superior temporal sulcus was associated with trustworthiness judgments, the fusiform gyrus was recruited by attractiveness judgments. In our study we found a volume increase of the fusiform gyrus for participants of sports teams and social clubs, but none for participants of religious group. With respect to the posterior superior temporal sulcus we found volume decrease within the group of sports team attendees and volume increase within attendees of religious groups. This suggests that processing of facial properties of other individuals may be of greater importance in social clubs and sport teams than in religious groups.

Those regions with the most pronounced volume deviations in our study have been widely connected to characteristics of social participation such as social network size. A previous functional neuroimaging study with resting-state functional magnetic resonance imaging found a positive association between social network size and the connectivity strength between the amygdala and superior temporal sulcus, as well as that between fusiform gyrus and to vmPFC respectively (Bickart et al., 2012). A related study by Bickert and colleagues (2011) linked the human social network size to brain structure of caudal inferior temporal sulcus, medial frontal cortex and ACC. The number of social contacts within one’s online social network (Facebook) has previously been linked to cortical volume deviation in the posterior superior temporal sulcus, as well as middle temporal gyrus (Kanai et al., 2012). Extending these results from human to non-human primates, macaques with a larger social network showed an increase in grey matter volume in rostral prefrontal cortex, ACC and superior temporal sulcus (Sallet et al., 2011). Our analysis regarding social participation highlighted several regions that were mentioned in the context of social network size. This indicates, that social participation is linked to a larger social network.

Only recently, the relevance of the DMN in the context of a lack of social connection, in other words loneliness has been highlighted (Spreng et al., 2020). Based on the results in Spreng and colleagues (2020) and those of our study, we can say that overlapping networks are implicated both for social isolation and social participation. Results included similar volume deviations in regions for participants feeling lonely, such ACC, posterior superior temporal sulcus and fusiform gyrus. Even on the network level, the relevance of DMN within lonely participants was alike the results within the groups of social participation. Hence, loneliness and social participation seem to tap on similar brain networks and may even be viewed as opposite ends of a continuum. Besides loneliness as a subjective form of social isolation, social support reflects an objective form of social isolation.

Social support is another key construct closely related to social participation. The construct of social support comprises closest friends and family, with a direct impact on health and well-being (Dunbar, 2018). Perhaps counterintuitive at first glance, results for social support and variance in brain regions showed less concordance to our results for social participation (Schurz et al., 2021). Although both assess lifestyle phenotypes that are related to forms of social cognition, only the effect of the limbic network and the volume deviation in the ACC appear to be more closely borne out by our present results. We speculate that the relevance of DMN and limbic system for social participation might derive from two sides. On the one hand, subjective aspects may be rooted in the DMN and reflect components like the sense of belonging to a group, hence less loneliness. Similarly, objective aspects of social participation may be linked to the limbic system and are possibly referring to a larger social network, providing social support.

On the conceptual level, social participation is not only used in various ways as a term but also measured in sometimes diverging ways (Chang & Coster, 2014), with constituent aspects consisting of a social role (e.g., daughter/son, friend, club member) and a social task (e.g., work environment, school). To date, the construct of social participation is predominantly discussed in the context of rehabilitation and healthy aging, being extensively investigated as an outcome measure (Douglas et al., 2017; Piškur et al., 2014).

As an empirical approach taken by our study, we examined three concrete forms of structured social participation with regular attendance. Sharing the same interest might in part be a consequence of shared personality traits, potentially linked to corresponding neural correlates. In support of this hypotheses, similarity of (autistic) personality traits in friendships of healthy adults was recently shown to be linked to friendship quality (Bolis et al., 2021). Especially as interindividual similarity in personality, social cognition and behaviour promote getting “in sync” and building meaningful social relationships (Hyon et al., 2020; Redcay & Schilbach, 2019).

In addition to a number of commonalities in brain structure and function, we also found differences between participants of sports teams, social clubs and religious groups. These differences might depend on the types of regular experienced interaction with the people who tend to be at the periphery of one’s social circles. For regular physical activity, positive effects on brain development and cognition in adolescence are presented (Herting & Chu, 2017). Herein reviewed neuroimaging studies reported volume increase in hippocampus and lingual gyrus was related to aerobic training. Improved cognition due to physical activity included executive functions, cognitive flexibility and inhibitory control. A review study attributes an increase in grey matter volume to physical activity for all brain regions except for superior temporal gyrus and fusiform gyrus (Batouli & Saba, 2017). However, in our study we found volume deviation for these two regions in sport team members as well. This in fact may be driven by social aspects of sport participation rather than physical activity.

For groups related to religious beliefs and spirituality, altered functional coupling patterns in the DMN have been reported and was discussed in the context of mystical and “insight” experiences (van Elk & Aleman, 2017). Differences in social processing between religious and nonreligious participants were found but controversial discussed at the same time, with special attention to peer influence and membership in religious groups (Grafman et al., 2020). Furthermore, no consistent grey matter volume differences for religiosity and mystical experiences were found in a recent voxel-based morphometry study (van Elk & Snoek, 2020). A meta-analytic study found no positive longevity effects based on individuals’ beliefs but still suggests positive health effects from simply belonging to a religious group (Shor & Roelfs, 2013). These effects include health-related behaviours and a sense of comfort and meaning, as the sheer result of social participation.

Adolescents with regular participation in different types of social clubs had a healthier lifestyle and protective effect of belonging to at least one club was described (Borraccino et al., 2020; Zambon et al., 2010). Older adults participating in social activities benefit from reduction of stress and improvements in several important dimensions of mental health (Mackenzie & Abdulrazaq, 2021). Another systematic review associated long-term commitment to social activity groups with executive functioning, social network size and global cognition (e.g. memory, executive function; Kelly et al., 2017). The surveyed intervention studies included sport groups (e.g., Tai Chi, walking) and social clubs (e.g., photo group, quilt group).

Hence, societies offering more ample opportunities for participation in sports, religious or social activities, not only dampen loneliness and cognitive decline but improve people’s resilience and might reduce the risk for dementia. In contrast, loneliness and social isolation are associated with loss of cognitive capacity (Lara et al., 2019), such as measured by Wechsler Adult Intelligence Scale, Consortium to Establish a Registry for Alzheimer’s Disease and digit span test. The previous studies provide various results concerning the impact of the particular nature of the investigated groups. Reported positive effects on health and cognition as well as deviations in brain structures might primarily derive from the recurring engagement with a group per se. In our study, we reported minor differences in brain structure and function among all three examined groups of social participation. However, similarities tended to dominate on structural and functional level in our collective findings.

Within our results, convergence was also shown across the different demographic profiling analyses. These brain-phenotype associations showed high similarity within most factors among participants of sports teams, religious groups and social clubs. Highest overlap across all three groups was observed in the factors television consumption, number of siblings, health-related lifestyle behaviour and psychological conditions. Although the direction of the revealed associations has no single answer in the fully multivariate setting, previous research reported general positive effects of group participation on substance use (Elder et al., 2000). Indeed, a wide range of previous studies presented heterogenous effects of physical activity on increased risk for substance use among adolescents (Murray et al., 2021; Terry-Mcelrath et al., 2011). While the similarity within psychological conditions, health factors and substance-use (limited to tobacco, nicotine, alcohol) is in line with recent findings in people receiving high social support (Schurz et al., 2021), being explained by general health effects of social relations (Holt-Lunstad et al., 2010), watching television and family size stand out. The individual consumption of television of adolescents was associated with the TV consumption of their peer group (Fletcher, 2006). Furthermore, siblings foster social competence, especially in a young age (Downey et al., 2015). More specific, perspective taking can be improved by siblings but findings vary and depend on the family context (Sang & Nelson, 2017). Again, despite of differences in the participated activity, commonalities exceed among the investigated groups in the demographic profiling analyses.

Membership to social groups scaffolds human life in society. Extending previous experimental neuroscience evidence, our investigation shows that brain substrates of social participation are interrelated with health-related concepts like social support and psychological well-being at the population level. Among all three examined types of groups, we identified the DMN and limbic network as central for social participation. Both highlighted networks gained further relevance in the context of belonging to a group, as aspects of everyday life participation were studied in a population cohort and could be related to additional demographic and everyday-life information. In a comprehensive demographic profiling analysis, we here find concrete benefits of social participation in groups such as reduced substance use and improved psychological well-being.

Overall, our collective findings could be taken to suggest that the concrete type of social participation may be of less importance than the regular attendance itself. Looking for a way to harness the positive effects of social participation, this calls for accessible forms of routine interventions in cohesive social groups, sometimes described as ‘social prescribing’. This is all the more important in periods of social isolation in which a lack of social participation takes its toll on mental health.

## Acknowledgements

This project has been made possible by the Brain Canada Foundation, through the Canada Brain Research Fund, as well as by NIH grant R01AG068563A and the Canadian Institutes of Health Research. DB was also supported by the Healthy Brains Healthy Lives initiative (Canada First Research Excellence fund), and by the CIFAR Artificial Intelligence Chairs program (Canada Institute for Advanced Research), as well as Research Award and Teaching Award by Google.

## References

Adams, K. B., Leibbrandt, S., & Moon, H. (2011). A critical review of the literature on social and leisure activity and wellbeing in later life. Ageing and Society, 31(4), 683–712. https://doi.org/10.1017/S0144686X10001091

Alfaro-Almagro, F., Jenkinson, M., Bangerter, N. K., Andersson, J. L. R., Griffanti, L., Douaud, G., Sotiropoulos, S. N., Jbabdi, S., Hernandez-Fernandez, M., Vallee, E., Vidaurre, D., Webster, M., McCarthy, P., Rorden, C., Daducci, A., Alexander, D. C., Zhang, H., Dragonu, I., Matthews, P. M.,… Smith, S. M. (2018). Image processing and Quality Control for the first 10,000 brain imaging datasets from UK Biobank. NeuroImage, 166, 400–424. https://doi.org/10.1016/j.neuroimage.2017.10.034

Amft, M., Bzdok, D., Laird, A., Fox, P. T., Schilbach, L., & Eickhoff, S. B. (2015). Definition and characterization of an extended social-affective default network. In Brain Struct Funct (Vol. 220, Issue 2). https://doi.org/10.1007/springerreference_184020

Ammar, A., Chtourou, H., Boukhris, O., Trabelsi, K., Masmoudi, L., Brach, M., Bouaziz, B., Bentlage, E., How, D., Ahmed, M., Mueller, P., Mueller, N., Hsouna, H., Aloui, A., Hammouda, O., Paineiras-Domingos, L. L., Braakman-Jansen, A., Wrede, C., Bastoni, S.,… Hoekelmann, A. (2020). Covid-19 home confinement negatively impacts social participation and life satisfaction: A worldwide multicenter study. International Journal of Environmental Research and Public Health, 17(6237), 1–17. https://doi.org/10.3390/ijerph17176237

Amodio, D. M., & Frith, C. D. (2006). Meeting of minds: The medial frontal cortex and social cognition. Nature Reviews Neuroscience, 7(4), 268–277. https://doi.org/10.1038/nrn1884

Andersson, J., Jenkinson, M., & Smith, S. (2007). Non-linear registration aka Spatial normalisation FMRIB Technical Report TR07JA2. FMRIB. Analysis Group of the University of Oxford.

Atroszko, P., Baginska, P., Mokosinska, M., Sawicki, A., & Atroszko, B. (2015). Validity and reliability of single-item self-report measures of general quality of life, general health and sleep quality. CER Compar Eur Res, 216.

Batouli, S. A. H., & Saba, V. (2017). At least eighty percent of brain grey matter is modifiable by physical activity: A review study. Behavioural Brain Research, 332(May), 204–217. https://doi.org/10.1016/j.bbr.2017.06.002

Beckmann, C., & Smith, S. (2004). Probabilistic independent component analysis for functional magnetic resonance imaging. IEEE Trans Med Imaging, 23, 137–152.

Bickart, K. C., Hollenbeck, M. C., Barrett, L. F., & Dickerson, B. C. (2012). Intrinsic amygdala-cortical functional connectivity predicts social network size in humans. Journal of Neuroscience, 32(42), 14729–14741. https://doi.org/10.1523/JNEUROSCI.1599-12.2012

Bickart, K. C., Wright, C. I., Dautoff, R. J., Dickerson, B. C., & Barrett, L. F. (2011). Amygdala volume and social network size in humans. Nature Neuroscience, 14(2), 163–164. https://doi.org/10.1038/nn.2724

Bolis, D., Lahnakoski, J. M., Seidel, D., Tamm, J., & Schilbach, L. (2021). Interpersonal similarity of autistic traits predicts friendships quality. Social Cognitive and Affective Neuroscience, 16(1-2), 222–231.

Borraccino, A., Lazzeri, G., Kakaa, O., Bad’ura, P., Bottigliengo, D., Dalmasso, P., & Lemma, P. (2020). The contribution of organised leisure-time activities in shaping positive community health practices among 13-and 15-year-old adolescents: Results from the health behaviours in school-aged children study in italy. International Journal of Environmental Research and Public Health, 17(18), 1–12. https://doi.org/10.3390/ijerph17186637

Boyce, T., & Brown, C. (2017). Engagement and participation for health equity. In World Health Organization Europe. http://www.euro.who.int/_data/assets/pdf_file/0005/353066/Engagement-and-Participation-HealthEquity.pdf?ua=1

Bzdok, D., Langner, R., Caspers, S., Kurth, F., Habel, U., Zilles, K., Laird, A., & Eickhoff, S. B. (2011). ALE meta-analysis on facial judgments of trustworthiness and attractiveness. Brain Structure and Function, 215(3–4), 209–223. https://doi.org/10.1007/s00429-010-0287-4

Bzdok, D, Eickenberg, M., Varoquaux, G., & Thirion, B. (2017). Hierarchical region-network sparsity for high-dimensional inference in brain imaging. Inf Process Med Imaging, 10265, 323–335.

Bzdok, Danilo, & Dunbar, R. I. M. (2020). The Neurobiology of Social Distance. Trends in Cognitive Sciences, 24(9), 717–733. https://doi.org/10.1016/j.tics.2020.05.016

Bzdok, Danilo, Floris, D. L., & Marquand, A. F. (2020). Analysing brain networks in population neuroscience: A case for the Bayesian philosophy. Philosophical Transactions of the Royal Society B: Biological Sciences, 375(1796), 1–11. https://doi.org/10.1098/rstb.2019.0661

Bzdok, Danilo, Langner, R., Hoffstaedter, F., Turetsky, B. I., Zilles, K., & Eickhoff, S. B. (2012). The modular neuroarchitecture of social judgments on faces. Cerebral Cortex, 22(4), 951–961. https://doi.org/10.1093/cercor/bhr166

Chang, F. H., & Coster, W. J. (2014). Conceptualizing the construct of participation in adults with disabilities. Archives of Physical Medicine and Rehabilitation, 95(9), 1791–1798. https://doi.org/10.1016/j.apmr.2014.05.008

Che, X. W., Wei, D. T., Li, W. F., Li, H. J., Qiao, L., Qiu, J., Zhang, Q. L., & Liu, Y. J. (2014). The correlation between gray matter volume and perceived social support: A voxel-based morphometry study. Social Neuroscience, 9(2), 152–159. https://doi.org/10.1080/17470919.2013.873078

Chen, A. C., Welsh, R. C., Liberzon, I., & Taylor, S. F. (2010). “Do I like this person?” A network analysis of midline cortex during a social preference task. NeuroImage, 51(2), 930–939. https://doi.org/10.1016/j.neuroimage.2010.02.044

Chiao, C., Weng, L. J., & Botticello, A. L. (2011). Social participation reduces depressive symptoms among older adults: An 18-year longitudinal analysis in Taiwan. BMC Public Health, 11. https://doi.org/10.1186/1471-2458-11-292

Cicognani, E., Pirini, C., Keyes, C., Joshanloo, M., Rostami, R., & Nosratabadi, M. (2008). Social participation, sense of community and social well being: A study on American, Italian and Iranian University students. Social Indicators Research, 89(1), 97–112. https://doi.org/10.1007/s11205-007-9222-3

Cohen, S., & Hoberman, H. M. (1983). Positive Events and Social Supports as Buffers of Life Change Stress. Journal of Applied Social Psychology, 13(2), 99–125.

Cornwell, E. Y., & Waite, L. J. (2009). Social disconnectedness, perceived isolation, and health among older adults. Journal of Health and Social Behavior, 50(1), 31–48. https://doi.org/10.1177/002214650905000103

Cyranowski, J. M., Zill, N., Bode, R., Butt, Z., Kelly, M., Pilkonis, P., Salsman, J., & Cella, D. (2013). Assessing Social Support, Companionship, and Distress: NIH Toolbox Adult Social Relationship Scales. Health Psychol, 32(3), 293–301. https://doi.org/10.1037/a0028586.Assessing

Desikan, R., Ségonne, F., Fischl, B., Quinn, B., Dickerson, B., Blacker, D., Buckner, R., Dale, A., Maguire, R., & Hyman, BT, et al. (2006). An automated labeling system for subdividing the human cerebral cortex on MRI scans into gyral based regions of interest. NEUROIMAGE, 31, 968–980.

Dollinger, S., & Malmquist, D. (2009). Reliability and validity of single-item self-reports: With special relevance to college students’ alcohol use, religiosity, study, and social life. The Journal of General Psychology, 136(3), 231–241.

Douglas, H., Georgiou, A., & Westbrook, J. (2017). Social participation as an indicator of successful aging: An overview of concepts and their associations with health. Australian Health Review, 41(4), 455–462. https://doi.org/10.1071/AH16038

Downey, D. B., Condron, D. J., & Yucel, D. (2015). Number of Siblings and Social Skills Revisited Among American Fifth Graders. Journal of Family Issues, 36(2), 273–296. https://doi.org/10.1177/0192513X13507569

Dunbar, Robin I.M. (2018). The Anatomy of Friendship. Trends in Cognitive Sciences, 22(1), 32–51.

Efron, B. (2012). Large-scale inference: empirical bayes methods for estimation, testing, and prediction (p. Cambridge (UK): Cambridge University Press.).

Efron, B., & Tibshirani, R. (1994). An introduction to the bootstrap.

Elder, C., Leaver-Dunn, D., Wang, M. Q., Nagy, S., & Green, L. (2000). Organized Group Activity as a Protective Factor Against Adolescent Substance Use. American Journal of Health Behavior, 24(2), 108–113.

Feldman, R. (2020). What is resilience: an affiliative neuroscience approach. World Psychiatry, 19(2), 132–150. https://doi.org/10.1002/wps.20729

Finlayson-Short, L., Davey, C. G., & Harrison, B. J. (2020). Neural correlates of integrated self and social processing. Social Cognitive and Affective Neuroscience, 941–949.

Fletcher, J. (2006). Social interactions in adolescent television viewing. Archives of Pediatrics and Adolescent Medicine, 160(4), 383–386. https://doi.org/10.1001/archpedi.160.4.383

Francés, F., La Parra, D., Martínez Asunción, M. R., Ortiz Barreda, G., & Briones Vozmediano, E. (2016). Toolkit on social participation. In World Health Organization Europe. https://www.euro.who.int/__data/assets/pdf_file/0003/307452/Toolkit-social-partecipation.pdf

Frith, U., & Frith, C. (2010). The social brain: Allowing humans to boldly go where no other species has been. Philosophical Transactions of the Royal Society B: Biological Sciences, 365(1537), 165–175. https://doi.org/10.1098/rstb.2009.0160

Gan, G., Ma, R., Reichert, M., Giurgiu, M., Ebner-Priemer, U. W., Meyer-Lindenberg, A., & Tost, H. (2021). Neural Correlates of Affective Benefit From Real-life Social Contact and Implications for Psychiatric Resilience. JAMA Psychiatry, 5–7. https://doi.org/10.1001/jamapsychiatry.2021.0560

Gelman, A., Carlin, J., Stern, H., & Rubin, D. (2014). Bayesian data analysis. Vol 2.

Grafman, J., Cristofori, I., Zhong, W., & Bulbulia, J. (2020). The Neural Basis of Religious Cognition. Current Directions in Psychological Science, 29(2), 126–133. https://doi.org/10.1177/0963721419898183

Griffanti, L., Salimi-Khorshidi, G., Beckmann, C., Auerbach, E., Douaud, G., Sexton, C., E, Z., Ebmeier, K., Filippini, N., & Mackay, C. (2014). ICA-based artefact removal and accelerated fMRI acquisition for improved resting state network imaging. Neuroimage, 95, 232–247.

Hartwigsen, G., Bengio, Y., & Bzdok, D. (2021). How does hemispheric specialization contribute to human-defining cognition? Neuron, 1–16. https://doi.org/10.1016/j.neuron.2021.04.024

Hawkley, L. C., Browne, M. W., & Cacioppo, J. T. (2005). How can I connect with thee? Let me count the ways. Psychological Science, 16(10), 798–804. https://doi.org/10.1111/j.1467-9280.2005.01617.x

Hawkley, L. C., Burleson, M. H., Berntson, G. G., & Cacioppo, J. T. (2003). Loneliness in Everyday Life: Cardiovascular Activity, Psychosocial Context, and Health Behaviors. Journal of Personality and Social Psychology, 85(1), 105–120. https://doi.org/10.1037/0022-3514.85.1.105

Herting, M. M., & Chu, X. (2017). Exercise, cognition, and the adolescent brain. Birth Defects Research, 109(20), 1672–1679. https://doi.org/10.1002/bdr2.1178

Hiser, J., & Koenigs, M. (2018). The multifaceted role of ventromedial prefrontal cortex in emotion, decision-making, social cognition, and psychopathology. Biol Psychiatry, 83(8), 638–647. https://doi.org/10.1016/j.biopsych.2017.10.030.The

Holt-Lunstad, J., Smith, T. B., & Layton, J. B. (2010). Social relationships and mortality risk: A meta-analytic review. PLoS Medicine, 7(7). https://doi.org/10.1371/journal.pmed.1000316

Hyatt, C. J., Calhoun, V. D., Pearlson, G. D., & Assaf, M. (2015). Specific default mode subnetworks support mentalizing as revealed through opposing network recruitment by social and semantic FMRI tasks. Human Brain Mapping, 36(8), 3047–3063. https://doi.org/10.1002/hbm.22827

Hyon, R., Youm, Y., Kim, J., Chey, J., Kwak, S., & Parkinson, C. (2020). Similarity in functional brain connectivity at rest predicts interpersonal closeness in the social network of an entire village. Proceedings of the National Academy of Sciences of the United States of America, 117(52), 33149–33160. https://doi.org/10.1073/PNAS.2013606117

Isik, L., Koldewyn, K., Beeler, D., & Kanwisher, N. (2017). Perceiving social interactions in the posterior superior temporal sulcus. Proceedings of the National Academy of Sciences of the United States of America, 114(1), E9145–E9152. https://doi.org/10.1073/pnas.1721071115

Jenkinson, M., Bannister, P., Brady, M., & Smith, S. (2002). Improved optimization for the robust and accurate linear registration and motion motion correction of brain images. NEUROIMAGE, 17, 825–841.

Jenkinson, M., & Smith, S. (2001). A global optimisation method for robust affine registration of brain images. Med Image Anal, 5, 143–156.

Kanai, R., Bahrami, B., Roylance, R., & Rees, G. (2012). Online social network size is reflected in human brain structure. Proceedings of the Royal Society B: Biological Sciences, 279, 1327–1334. https://doi.org/10.1098/rspb.2011.1959

Kelly, M. E., Duff, H., Kelly, S., McHugh Power, J. E., Brennan, S., Lawlor, B. A., & Loughrey, D. G. (2017). The impact ofsocial activities, social networks, social support and social relationships on the cognitive functioning of healthy older adults: A systematic review. Systematic Reviews, 6(259). https://doi.org/10.1186/s13643-017-0632-2

Kiesow, H., Dunbar, R., Kable, J., Kalenscher, T., Vogeley, K., Schilbach, L., Marquand, A., Wiecki, T., & Bzdok, D. (2020). 10,000 social brains: sex differentiation inhuman brain anatomy. Sci Adv, 6(eaaz1170), 1–12.

Lamblin, M., Murawski, C., Whittle, S., & Fornito, A. (2017). Social connectedness, mental health and the adolescent brain. Neuroscience and Biobehavioral Reviews, 80, 57–68. https://doi.org/10.1016/j.neubiorev.2017.05.010

Lara, E., Caballero, F. F., Rico-Uribe, L. A., Olaya, B., Haro, J. M., Ayuso-Mateos, J. L., & Miret, M. (2019). Are loneliness and social isolation associated with cognitive decline? International Journal of Geriatric Psychiatry, 34(11), 1613–1622. https://doi.org/10.1002/gps.5174

Law, M. (2002). Participation in the occupations of everyday life. American Journal of Occupational Therapy, 56(6), 640–649. https://doi.org/10.5014/ajot.56.6.640

Lee, T.-C. (2014). Trilogy of body imaginary: Dance/movement therapy for a psychiatric patient with depression. Arts in Psychotherapy, 41(4), 400–408. https://doi.org/10.1016/j.aip.2014.07.006

Leone, T., & Hessel, P. (2016). The effect of social participation on the subjective and objective health status of the over-fifties: Evidence from SHARE. Ageing and Society, 36(5), 968–987. https://doi.org/10.1017/S0144686X15000148

Leopold, A., Krueger, F., Dal monte, O., Pardini, M., Pulaski, S. J., Solomon, J., & Grafman, J. (2012). Damage to the left ventromedial prefrontal cortex impacts affective theory of mind. Social Cognitive and Affective Neuroscience, 7(8), 871–880. https://doi.org/10.1093/scan/nsr071

Lestari, S. K., de Luna, X., Eriksson, M., Malmberg, G., & Ng, N. (2021). A longitudinal study on social support, social participation, and older Europeans’ Quality of life. SSM - Population Health, 13(February). https://doi.org/10.1016/j.ssmph.2021.100747

Levasseur, M., Desrosiers, J., & Whiteneck, G. (2010). Accomplishment level and satisfaction with social participation of older adults: Association with quality of life and best correlates. Quality of Life Research, 19(5), 665–675. https://doi.org/10.1007/s11136-010-9633-5

Levasseur, M., Richard, L., Gauvin, L., & Raymond, É. (2010). Inventory and analysis of definitions of social participation found in the aging literature: Proposed taxonomy of social activities. Social Science and Medicine, 71(12), 2141–2149. https://doi.org/10.1016/j.socscimed.2010.09.041

Lewis, P. A., Rezaie, R., Brown, R., Roberts, N., & Dunbar, R. I. M. (2011). Ventromedial prefrontal volume predicts understanding of others and social network size. NeuroImage, 57(4), 1624–1629. https://doi.org/10.1016/j.neuroimage.2011.05.030

Luhmann, M., & Hawkley, L. C. (2016). Age Differences in Loneliness from Late Adolescence to Oldest Old Age. Dev Psychol, 52(6), 943–959. https://doi.org/10.1037/dev0000117.Age

Mackenzie, C. S., & Abdulrazaq, S. (2021). Social engagement mediates the relationship between participation in social activities and psychological distress among older adults. Aging & Mental Health, 25(2), 299–305.

Maier, S. F., & Watkins, L. R. (2010). Role of the medial prefrontal cortex in coping and resilience. Brain Research, 1355, 52–60. https://doi.org/10.1016/j.brainres.2010.08.039

Maij, D. L. R., Van Harreveld, F., Gervais, W., Schrag, Y., Mohr, C., & Van Elk, M. (2017). Mentalizing skills do not differentiate believers from non-believers, but credibility enhancing displays do. In PLoS ONE (Vol. 12, Issue 8). https://doi.org/10.1371/journal.pone.0182764

Mars, R. B., Neubert, F. X., Noonan, M. A. P., Sallet, J., Toni, I., & Rushworth, M. F. S. (2012). On the relationship between the “default mode network” and the “social brain.” Frontiers in Human Neuroscience, 6(JUNE 2012), 1–9. https://doi.org/10.3389/fnhum.2012.00189

Mashek, D., Cannaday, L., & Tangney, J. (2007). Inclusion of community in self scale: a single-item pictorial measure of community connectedness. Journal of Community Psychology, 35(2), 257–275. https://doi.org/10.1002/jcop

Miller, K. L., Alfaro-Almagro, F., Bangerter, N. K., Thomas, D. L., Yacoub, E., Xu, J., Bartsch, A. J., Jbabdi, S., Sotiropoulos, S. N., Andersson, J. L. R., Griffanti, L., Douaud, G., Okell, T. W., Weale, P., Dragonu, I., Garratt, S., Hudson, S., Collins, R., Jenkinson, M.,… Smith, S. M. (2016). Multimodal population brain imaging in the UK Biobank prospective epidemiological study. Nature Neuroscience, 19(11), 1523–1536. https://doi.org/10.1038/nn.4393

Morita, T., Asada, M., & Naito, E. (2021). Gray-matter expansion of social brain networks in individuals high in public self-consciousness. Brain Sciences, 11(374). https://doi.org/10.3390/brainsci11030374

Murray, R. M., Sabiston, C. M., Doré, I., Bélanger, M., & O’Loughlin, J. L. (2021). Longitudinal associations between team sport participation and substance use in adolescents and young adults. Addictive Behaviors, 116, 106798.

Mwilambwe-Tshilobo, L., Ge, T., Chong, M., Ferguson, M. A., Misic, B., Burrow, A. L., Leahy, R. M., & Spreng, R. N. (2019). Loneliness and meaning in life are reflected in the intrinsic network architecture of the brain. Social Cognitive and Affective Neuroscience, 14(4), 423–433. https://doi.org/10.1093/scan/nsz021

Ong, A. D., Uchino, B. N., & Wethington, E. (2016). Loneliness and Health in Older Adults: A Mini-Review and Synthesis. Gerontology, 62(4), 443–449. https://doi.org/10.1159/000441651

Oosterhof, N. N., & Todorov, A. (2008). The functional basis of face evaluation. Proceedings of the National Academy of Sciences of the United States of America, 105(32), 11087–11092. https://doi.org/10.1073/pnas.0805664105

Piškur, B., Daniëls, R., Jongmans, M. J., Ketelaar, M., Smeets, R. J. E. M., Norton, M., & Beurskens, A. J. H. M. (2014). Participation and social participation: Are they distinct concepts? Clinical Rehabilitation, 28(3), 211–220. https://doi.org/10.1177/0269215513499029

Powell, J., Lewis, P. A., Roberts, N., García-Fiñana, M., & Dunbar, R. I. M. (2012). Orbital prefrontal cortex volume predicts social network size: An imaging study of individual differences in humans. Proceedings of the Royal Society B: Biological Sciences, 279(1736), 2157–2162. https://doi.org/10.1098/rspb.2011.2574

Quinn, T., Adger, W. N., Butler, C., & Walker-Springett, K. (2020). Community resilience and well-being: an exploration of relationality and belonging after disasters. Annals of the American Association of Geographers, 111(2), 577–590.

Redcay, E., & Schilbach, L. (2019). Using second-person neuroscience to elucidate the mechanisms of social interaction. 20(August), 495–505.

Sallet, J., Mars, R. B., Noonan, M. P., Andersson, J. L., O’Reilly, J. X., Jbabdi, S., Croxson, P. L., Jenkinson, M., Miller, K. L., & Rushworth, M. F. S. (2011). Social Network Size Affects Neural Circuits in Macaques. Science, 334(6056), 697–700. https://doi.org/10.4159/harvard.9780674333987.c22

Salvatier, J., Wiecki, T., & Fonnesbeck, C. (2016). Probabilistic programming in python using PyMC3. PeerJ Computer Science, 2(e55), 1–24.

Sang, S. A., & Nelson, J. A. (2017). The effect of siblings on children’s social skills and perspective taking. Infant and Child Development, 26(6), 1–10. https://doi.org/10.1002/icd.2023

Scarf, D., Moradi, S., McGaw, K., Hewitt, J., Hayhurst, J. G., Boyes, M., Ruffman, T., & Hunter, J. A. (2016). Somewhere I belong: Long-term increases in adolescents’ resilience are predicted by perceived belonging to the in-group. British Journal of Social Psychology, 55(3), 588–599. https://doi.org/10.1111/bjso.12151

Schaefer, A., Kong, R., Gordon, E. M., Laumann, T. O., Zuo, X.-N., Holmes, A. J., Eickhoff, S. B., & Yeo, B. T. T. (2018). Local-Global Parcellation of the Human Cerebral Cortex from Intrinsic Functional Connectivity MRI. Cerebral Cortex, 28(9), 3095–3114. https://doi.org/10.1093/cercor/bhx179

Schilbach, L., Bzdok, D., Timmermans, B., Fox, P. T., Laird, A. R., Vogeley, K., & Eickhoff, S. B. (2012). Introspective Minds: Using ALE meta-analyses to study commonalities in the neural correlates of emotional processing, social & unconstrained cognition. PLoS ONE, 7(2). https://doi.org/10.1371/journal.pone.0030920

Schilbach, L., Eickhoff, S. B., Rotarska-Jagiela, A., Fink, G. R., & Vogeley, K. (2008). Minds at rest? Social cognition as the default mode of cognizing and its putative relationship to the “default system” of the brain. Consciousness and Cognition, 17(2), 457–467. https://doi.org/10.1016/j.concog.2008.03.013

Schurz, M., Uddin, L. Q., Kanske, P., Lamm, C., Sallet, J., Bernhardt, B. C., Mars, R. B., & Bzdok, D. (2021). Variability in Brain Structure and Function Reflects Lack of Peer Support. Cerebral Cortex, 1–16. https://doi.org/10.1093/cercor/bhab109

Shamay-Tsoory, S. G., Tomer, R., Berger, B. D., & Aharon-Peretz, J. (2003). Characterization of empathy deficits following prefrontal brain damage: The role of the right ventromedial prefrontal cortex. Journal of Cognitive Neuroscience, 15(3), 324–337. https://doi.org/10.1162/089892903321593063

Sharifian, N., & Grühn, D. (2019). The Differential Impact of Social Participation and Social Support on Psychological Well-Being: Evidence From the Wisconsin Longitudinal Study. International Journal of Aging and Human Development, 88(2), 107–126. https://doi.org/10.1177/0091415018757213

Shor, E., & Roelfs, D. J. (2013). The longevity effects of religious and nonreligious participation: A meta-analysis and meta-regression. Journal for the Scientific Study of Religion, 52(1), 120–145. https://doi.org/10.1111/jssr.12006

Sinha, R., Lacadie, C. M., Constable, R. T., & Seo, D. (2016). Dynamic neural activity during stress signals resilient coping. Proceedings of the National Academy of Sciences of the United States of America, 113(31), 8837–8842. https://doi.org/10.1073/pnas.1600965113

Sirven, N., & Debrand, T. (2008). Social participation and healthy ageing: An international comparison using SHARE data. Social Science and Medicine, 67(12), 2017–2026. https://doi.org/10.1016/j.socscimed.2008.09.056

Smith, S. (2002). Fast robust automated brain extraction. Hum Brain Mapp, 17, 143–155.

Smith, S., Zhang, Y., Jenkinson, M., Chen, J., Matthews, P., Federico, A., & De Stefano, N. (2002). Accurate, robust, and automated longitudinal and cross-sectional brain change analysis. NEUROIMAGE, 17, 479–489.

Spreng, R. N., Dimas, E., Mwilambwe-tshilobo, L., Dagher, A., Koellinger, P., Nave, G., Ong, A., Kernbach, J. M., Wiecki, T. V, Ge, T., Li, Y., Holmes, A. J., Yeo, B. T. T., Turner, G. R., Dubar, R. I. M., & Bzdok, D. (2020). The default network of the human brain is associated with perceived social isolation. Nature Communications, 1–11. http://dx.doi.org/10.1038/s41467-020-20039-w

Taebi, A., Kiesow, H., Vogeley, K., Schilbach, L., Bernhardt, B. C., & Bzdok, D. (2020). Population variability in social brain morphology for social support, household size and friendship satisfaction. Social Cognitive and Affective Neuroscience, 15(6), 635–647. https://doi.org/10.1093/scan/nsaa075

Takeuchi, H., Taki, Y., Sassa, Y., Hashizume, H., Sekiguchi, A., Fukushima, A., & Kawashima, R. (2014). Regional gray matter volume is associated with empathizing and systemizing in young adults. PLoS ONE, 9(1). https://doi.org/10.1371/journal.pone.0084782

Talò, C., Mannarini, T., & Rochira, A. (2014). Sense of Community and Community Participation: A Meta-Analytic Review. Social Indicators Research, 117(1), 1–28. https://doi.org/10.1007/s11205-013-0347-2

Terry-Mcelrath, Y. M., O’Malley, P. M., & Johnston, L. D. (2011). Exercise and substance use among american youth, 1991-2009. American Journal of Preventive Medicine, 40(5), 530–540. https://doi.org/10.1016/j.amepre.2010.12.021

Thoits, P. A. (2011). Mechanisms linking social ties and support to physical and mental health. Journal of Health and Social Behavior, 52(2), 145–161. https://doi.org/10.1177/0022146510395592

van Elk, M., & Aleman, A. (2017). Brain mechanisms in religion and spirituality: An integrative predictive processing framework. Neuroscience and Biobehavioral Reviews, 73, 359–378. https://doi.org/10.1016/j.neubiorev.2016.12.031

van Elk, M., & Snoek, L. (2020). The relationship between individual differences in gray matter volume and religiosity and mystical experiences: A preregistered voxel-based morphometry study. European Journal of Neuroscience, 51(3), 850–865. https://doi.org/10.1111/ejn.14563

Walbrin, J., Downing, P., & Koldewyn, K. (2018). Neural responses to visually observed social interactions. Neuropsychologia, 112(February), 31–39. https://doi.org/10.1016/j.neuropsychologia.2018.02.023

Wang, S., Tepfer, L. J., Taren, A. A., & Smith, D. V. (2020). Functional parcellation of the default mode network: a large-scale meta-analysis. Scientific Reports, 10(1), 1–13. https://doi.org/10.1038/s41598-020-72317-8

WHO (World Health Organization). (2002). Active ageing: a policy framework. In Geneva, Switzerland: World Health Organization.

Zambon, A., Morgan, A., Vereecken, C., Colombini, S., Boyce, W., Mazur, J., Lemma, P., & Cavallo, F. (2010). The contribution of club participation to adolescent health: Evidence from six countries. Journal of Epidemiology and Community Health, 64(1), 89–95. https://doi.org/10.1136/jech.2009.088443

Zhang, Y., Brady, M., & Smith, S. (2001). Segmentation of brain MR images through a hidden Markov random field model and the expectation-maximization algorithm. IEEE Trans Med Imaging, 20, 45–57.

